# Bayesian selection of grammar productions for the language of thought

**DOI:** 10.1101/141358

**Authors:** S. Romano, A. Salles, M. Amalric, S. Dehaene, M. Sigman, S. Figueria

## Abstract

Probabilistic proposals of Language of Thoughts (LoTs) can explain learning across different domains as statistical inference over a compositionally structured hypothesis space. While frameworks may differ on how a LoT may be implemented computationally, they all share the property that they are built from a set of atomic symbols and rules by which these symbols can be combined. In this work we show how the set of productions of a LoT grammar can be effectively selected from a broad repertoire of possible productions by an inferential process starting from experimental data. We then test this method in the *language of geometry*, a specific LoT model (Amalric et al., 2017). Finally, despite the fact of the geometrical LoT not being a universal (i.e. Turing-complete) language, we show an empirical relation between a sequence’s *probability* and its *complexity* consistent with the theoretical relationship for universal languages described by Levin’s Coding Theorem.

---

> It was not only difficult for him to understand that the generic term dog embraced so many unlike specimens of differing sizes and different forms; he was disturbed by the fact that a dog at three-fourteen (seen in profile) should have the same name as the dog at three-fifteen (seen from the front). (…) With no effort he had learned English, French, Portuguese and Latin. I suspect, however, that he was not very capable of thought. To think is to forget differences, generalize, make abstractions. In the teeming world of Funes, there were only details, almost immediate in their presence. (Borges, 1944)

In his fantasy story, the writer Jorge Luis Borges described a fictional character, Funes, capable of remembering every detail of his life but not being able to generalize any of that data into mental categories and hence –Borges stressed– not capable of thinking.

Researchers have modeled these mental categories or conceptual classes with two classical approaches: in terms of similarity to a generic example or prototype (Nosofsky, 1986; Rosch, 1999; Rosch & Mervis, 1975; Rosch, Simpson, & Miller, 1976) or based on a symbolic/rule-like representation (Boole, 1854; Fodor, 1975; Gentner, 1983).

Symbolic approaches like the *language of thought* (LoT) hypothesis (Fodor, 1975), claim that thinking takes form in a sort of mental language, composed of a limited set of atomic symbols that can be combined to form more complex structures following combinatorial rules.

Despite criticisms and objections (Aydede, 1997; Blackburn, 1984; Knowles, 1998; Loewer & Rey, 1991), symbolic approaches—in general— and the LoT hypothesis —in particular— have gained some renewed attention with recent results that might explain learning across different domains as statistical inference over a compositionally structured hypothesis space (Piantadosi & Jacobs, 2016; Tenenbaum, Kemp, Griffiths, & Goodman, 2011).

The LoT is not necessarily unique. In fact, the form that it takes has been modeled in many different ways depending on the problem domain: numerical concept learning (Piantadosi, Tenenbaum, & Goodman, 2012), sequence learning (Amalric et al., 2017; Romano, Sigman, & Figueira, 2013; Yildirim & Jacobs, 2015), visual concept learning (Ellis, Solar-Lezama, & Tenenbaum, 2015), theory learning (Ullman, Goodman, & Tenenbaum, 2012), etc.

While frameworks may differ on how a LoT may be implemented computationally, they all share the property of being built from a set of atomic symbols and rules by which they can be combined to form new and more complex expressions.

Most studies of LoTs have focused on the compositional aspect of the language, which has either been modeled within a Bayesian (Tenenbaum et al., 2011) or a Minimum Description Length (MDL) framework (Amalric et al., 2017; Goldsmith, 2001, 2002; Romano et al., 2013).

The common method is to define a grammar with a set of productions based on operations that are intuitive to researchers and then study how different inference processes match regular patterns in human learning. A recent study by Piantadosi, Tenenbaum, and Goodman (2016) puts the focus on the process of how to empirically choose the set of productions and how different LoT definitions can create different patterns of learning. Here, we move along that direction but use Bayesian inference to individuate the LoT instead of comparing several of them by hand.

Broadly, our aim is to propose a method to select the set of atomic symbols in an inferential process by pruning and trimming from a broad repertoire. More precisely, we test whether Bayesian inference can be used to decide the proper set of productions in a LoT defined by a context free grammar. These productions are derived from the subjects’ experimental data. In order to do this, a researcher builds a broader language with two sets of productions: 1) those for which she has a strong prior conviction that they should be used in the cognitive task, and 2) other productions that could be used to structure the data and extract regularities even if she believes are not part of the human reasoning repertoire for the task. With the new broader language, she should then turn the context free grammar that defines it into a probabilistic context free grammar (PCFG) and use Bayesian analysis to infer the probability of each production in order to choose the set that best explains the data.

In the next section we formalize this procedure and then apply it on the *language of geometry* presented by Amalric et al. (2017) in a recent study about geometrical sequence learning. This LoT defines a language with some basic geometric instructions as the grammar productions and then models their composition within the MDL framework. Our method, however, can be applied to any LoT model that defines a grammar, independently of whether its compositional aspect is modeled using a Bayesian framework or a MDL approach.

Finally, even with the recent surge of popularity of Bayesian inference and MDL in cognitive science, there are –to the best of our knowledge– no practical attempts to close the gap between probabilistic and complexity approaches to LoT models.

The theory of computation, through Levin’s Coding Theorem (Levin, 1974), exposes a remarkable relationship between the *Kolmogorov complexity* of a sequence and its *universal probability*, largely used in algorithmic information theory. Although both metrics are actually non-computable and defined over a universal prefix Turing Machine, we can apply both ideas to other non-universal Turing Machines in the same way that the concept of complexity used in MDL can be computed for specific, non-universal languages.

In this work, we examine the extent to which this theoretical prediction for infinite sequences holds empirically for a specific LoT, the *language of geometry*. Although the inverse logarithmic relationship between both metrics is proved for universal languages in the Coding Theorem, testing this same property for a particular non-universal language shows that the language shares some interesting properties of general languages. This constitutes a first step towards a formal link between probability and complexity modeling frameworks for LoTs.

## Bayesian inference for LoT’s productions

The project of Bayesian analysis of the LoT models concept learning using Bayesian inference in a grammatically structured hypothesis space (Goodman, Tenenbaum, Feldman, & Griffiths, 2008). Each LoT proposal is usually formalized by a context free grammar *𝒢* that defines the valid programs that can be generated, like in any other programming language. A program is a derivation tree of *𝒢* that needs to be interpreted or executed according to a given semantics in order to get an actual description of the concept in the cognitive task at hand. Finally, a Bayesian inference process is defined in order to infer the distribution of valid programs in *𝒢* from the observed data.

As explained above, our aim is to derive the productions of *𝒢* from the data, instead of just conjecturing them using a priori knowledge about the task. Prior work on LoTs has fit probabilities of productions in a context free grammar using Bayesian inference, however, the focus has been put in integrating out the production probabilities to better predict the data without changing the grammar definition (Piantadosi et al., 2016). Here, we want to study if the inference process could let us decide which productions can be safely pruned from the grammar. We introduce a generic method that can be used on any grammar to select and test the proper set of productions. Instead of using a fixed grammar and adjusting the probabilities of the productions to predict the data, we use Bayesian inference to rule out productions with probability lower than a certain threshold. This allows the researcher to validate the adequacy of the productions she has chosen for the grammar or even define one that is broad enough to express different regularities and let the method select the best set for the observed data.

To infer the probability for each production based on the observed data, we need to add a vector of probabilities *θ* associated with each production in order to convert the context free grammar *𝒢* into a probabilistic context free grammar (PCFG) (Manning & Schütze, 1999).

Let *D* = (*d*_1_, *d*_2_, …, *d*_*n*_) denote the list of concepts produced by the subjects in an experiment. This means that each *d*_*i*_ is a concept produced by a subject in each trial. Then, *P*(*θ* | *D*), the posterior probability of the weights of each production after the observed data, can be calculated by marginalizing over the possible programs that compute *D*:

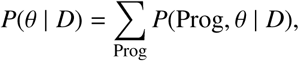

where each Prog = (*p*_1_, *p*_2_, …, *p*_*n*_) is a possible set of programs such that each *p*_*i*_ computes the corresponding concept *d*_*i*_.

We can use Bayesian inference to learn the corresponding programs Prog and the vector *θ* for each production in the grammar, applying Bayes rule in the following way:

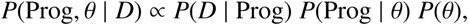

where *P*(*θ*) is a Dirichlet prior for *θ* and α its associated concentration vector hyper-parameter. The likelihood function can be calculated as follows:

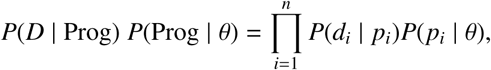

where *P*(*d*_*i*_ | *p*_*i*_) = 1 if the program *p*_*i*_ computes *d*_*i*_, and 0 otherwise, and 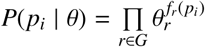 is the probability of the program *p*_*i*_ in the grammar, and *f*_*r*_ (*p*_*i*_) is the number of occurrences of the production *r* in program *p*_*i*_.

Calculating *P*(*θ* | *D*) directly is, however, not tractable since it requires to sum over all possible combinations of programs Prog for each of the possible values of *θ*. To this aim, then, we used a Gibbs Sampling (Geman & Geman, 1984) algorithm for PCFGs via Markov Chain Monte Carlo (MCMC) similar to the one proposed by Johnson, Griffiths, and Goldwater (2007), which alternates in each step of the chain between the two conditional distributions:

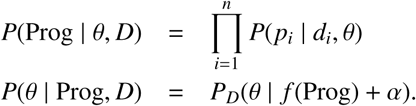

Here, *P*_*D*_ is the Dirichlet distribution where the positions of the vector α were updated by counting the occurrences of the corresponding productions for all programs *p*_*i*_ ∈ Prog.

In the next section, we apply this method to a specific LoT. We add a new set of ad-hoc productions to the grammar that can explain regularities but are not related to the cognitive task. If the method is effective, it should assign a low probability to the ad-hoc productions and instead favor the original set of productions selected by the researchers for the cognitive task.

This would not only provide empirical evidence about the adequacy of the choice of the original productions for the selected LoT but, more importantly, about the usefulness of Bayesian inference for selecting or testing the set of productions involved in different LoTs.

## The Language of Geometry: *𝒢eo*

The *language of geometry*, *𝒢eo*, is a probabilistic generator of sequences of movements on a regular octagon like the one in Figure 1.

**Figure 1.**
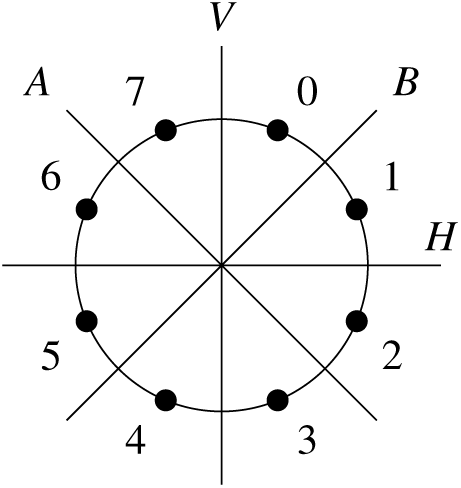
Σ points around a circle to map current position in the octagon, and the reflection axes.

The production rules of grammar *𝒢eo* were selected based on previous claims of the universality of certain human geometrical knowledge (Dehaene, Izard, Pica, & Spelke, 2006; Dillon, Huang, & Spelke, 2013; Izard, Pica, Dehaene, Hinchey, & Spelke, 2011) such as spatial notions (Landau, Gleitman, & Spelke, 1981; Lee, Sovrano, & Spelke, 2012) and detection of symmetries (Machilsen, Pauwels, & Wagemans, 2009; Westphal-Fitch, Huber, Gómez, & Fitch, 2012).

With these production rules, sequences are described by concatenating or repeating sequence of movements in the octagon. The original set of productions is shown in Table 1 and –besides the concatenation and repetition operators– it includes the following family of atomic geometrical transition productions: anticlockwise movements, staying at the same location, clockwise movements and symmetry movements.

**Table 1.**
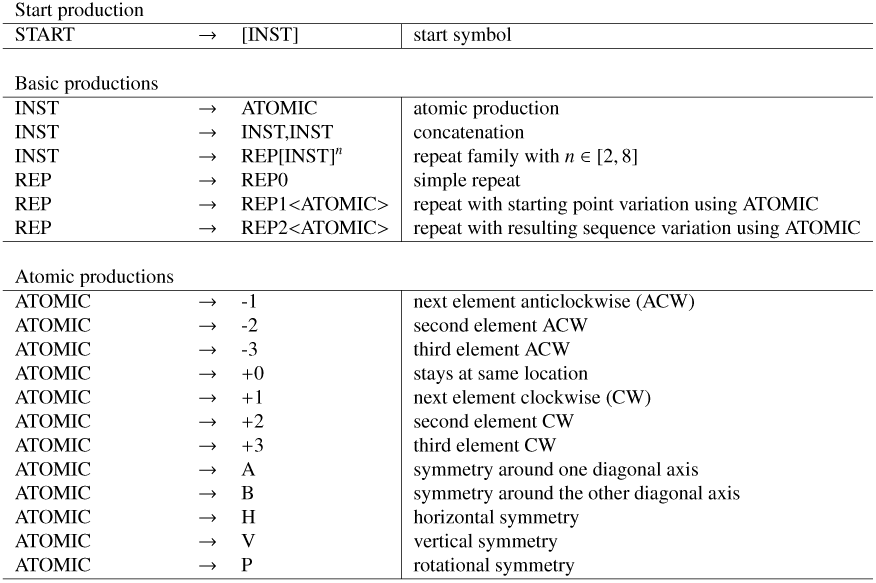
*Original Grammar*

The language actually supports not just a simple *n* times repetition of a block of productions, but it also supports two more complex productions in the repetition family: repeating with a change in the starting point after each cycle and repeating with a change to the resulting sequence after each cycle. More details about the formal syntax and semantics can be found in (Amalric et al., 2017), though they are not needed here.

Each program *p* generated by the grammar describes a mapping Σ → Σ^+^, for Σ = {0, …, 7}. Here, Σ^+^ represents the set of all (non empty) finite sequences over the alphabet Σ, which can be understood as a finite sequence of points in the octagon. These programs must then be executed or interpreted from a starting point in order to get the resulting sequence of points. Let *p* = [+1,+1] be a program, then *p*(0) is the result of executing *p* starting from point 0 (that is, sequence 1, 2) and *p*(4) is the result of executing the same program starting from point 4 in the octagon (sequence 5, 6)

Each sequence can be described with many different programs: from a simple concatenation of atomic productions to more compressed forms using repetitions. For example, to move through all the octagon clockwise one point at a time starting from point 0, one can use [+1,+1,+1,+1,+1,+1,+1,+1](0) or [REP[+1]^8^](0) or [REP[+1]^7^, +1](0), etc. To alternate 8 times between points 6 and 7, one can use a reflection production like [REP[A]^8^](6), or [REP[+1,-1]^4^](6).

## *𝒢eo*’s original experiment

To infer the productions from the observed data, we used the original data from the experiment in (Amalric et al., 2017). In the experiment, volunteers were exposed to a series of spatial sequences defined on an octagon and were asked to predict future locations. The sequences were selected according to their MDL in the *language of geometry* so that each sequence could be easily described with few productions.

### Participants

The data used in this work comes, except otherwise stated, from Experiment 1 in which participants were 23 French adults (12 female, mean age = 26.6, age range = 20 – 46) with college-level education. Data from Experiment 2 is later used when comparing adults and children results. In the later, participants where 24 preschoolers (minimal age = 5.33, max = 6.29, mean = 5.83 ± 0.05).

### Procedure

On each trial, the first two points from the sequence were flashed sequentially in the octagon and the user had to click on the next location. If the subject selected the correct location, she was asked to continue with the next point until the eight points of the sequences were completed. If there was an error at any point, the mistake was corrected, the sequence flashed again from the first point to the corrected point and the user asked to predict the next location. Each *d*_*i*_ ∈ Σ^8^ from our dataset *D* is thus the sequence of eight positions clicked in each subject’s trial. The detailed procedure can be found in the cited work.

## Extending *𝒢eo*’s grammar

We will now expand the original set of productions in G*eo* with a new set of productions that can also express regularities but are not related to any geometrical intuitions to test our Bayesian inference model.

In table 2 we show the new set of productions which includes instructions like moving to the point whose label is the square of the current location’s label, or using the current point location *i* to select the *i*^th^digit of a well-known number like *π* or Chaitin’s number^1^ (digits are returned in arithmetic module 8 to get a valid point for the next position). For example, PI(0) returns the first digit of π, that is PI(0) = 3 mod (8) = 3; and PI(1) = 1

**Table 2.**
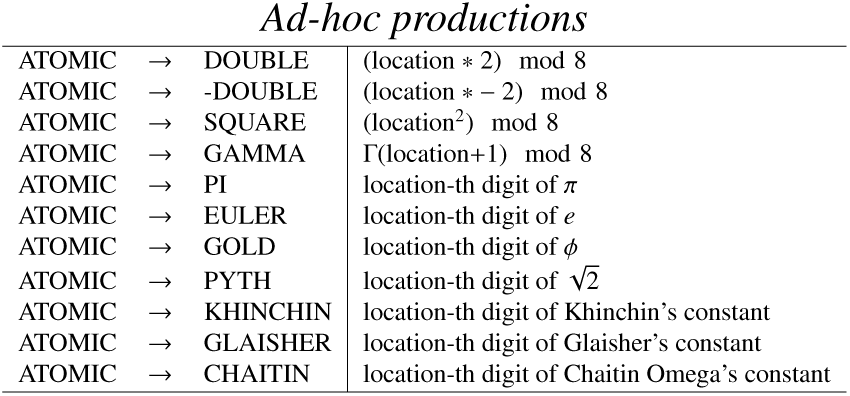
*Ad-hoc productions*

## Inference results for *𝒢eo*

To let the MCMC converge faster (and to later compare the concept’s probability with their corresponding MDL), we generated all the programs that explain each of the observed sequences from the experiment. In this way, we are able to sample from the exact distribution *P*(*p*_*i*_ | *d*_*i*_, *θ*) by sampling from a multinomial distribution of all the possible programs *p*_*i*_ that compute *d*_*i*_, where each *p*_*i*_ has probability of occurrence equal to *P*(*p*_*i*_ | *θ*).

To get an idea of the expressiveness of the grammar to generate different programs for a sequence and the cost of computing them, it is worth mentioning that there are more than 159 million programs that compute the 292 unique sequences generated by the subjects in the experiment, and that for each sequence there is an average of 546,713 programs (min = 10, 749, max = 5, 500, 026, σ = 693, 618).

Figure 2 shows the inferred *θ* for the observed sequences from subjects, with a unit concentration parameter for the Dirichlet prior, *α* = (1, …, 1). Each bar shows the mean probability and the standard error of each of the atomic productions after 50 steps of the MCMC, leaving the first 10 steps out as burn-in.

**Figure 2.**
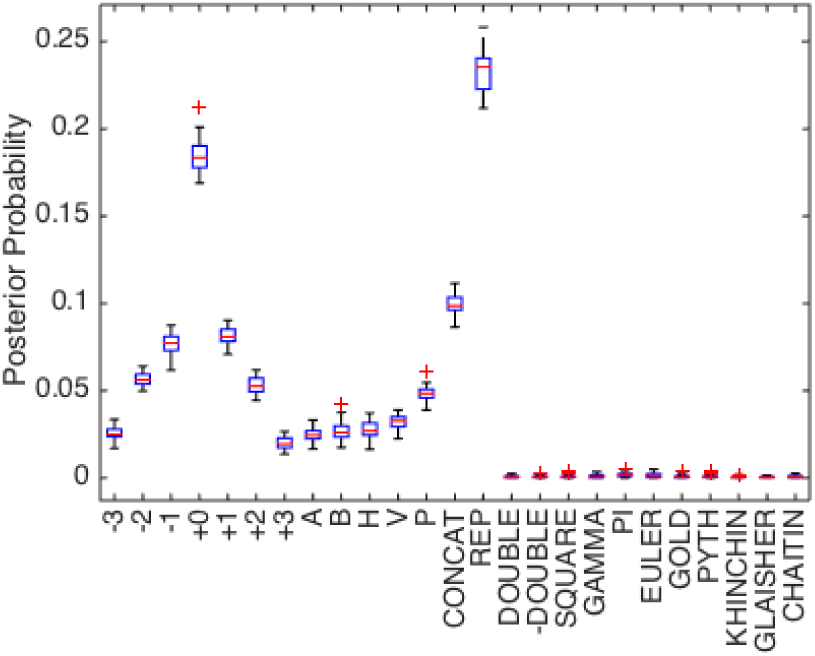
Inferred *θ*_*i*_ probability for each production in the grammar

Although 50 steps might seem low for a MCMC algorithm to converge, our method calculated *P*(*p*_*i*_ | *d*_*i*_, *θ*) exactly in order to speed up convergence and to be able to later compare the probability with the complexity from the original MDL model. In Figure 3, we show an example trace for four MCMC runs for *θ*_+0_, which corresponds to the atomic production +0, but is representative of the behavior of all *θ*_*i*_. (Figures for the full set of productions can be found in the Appendix).

**Figure 3.**
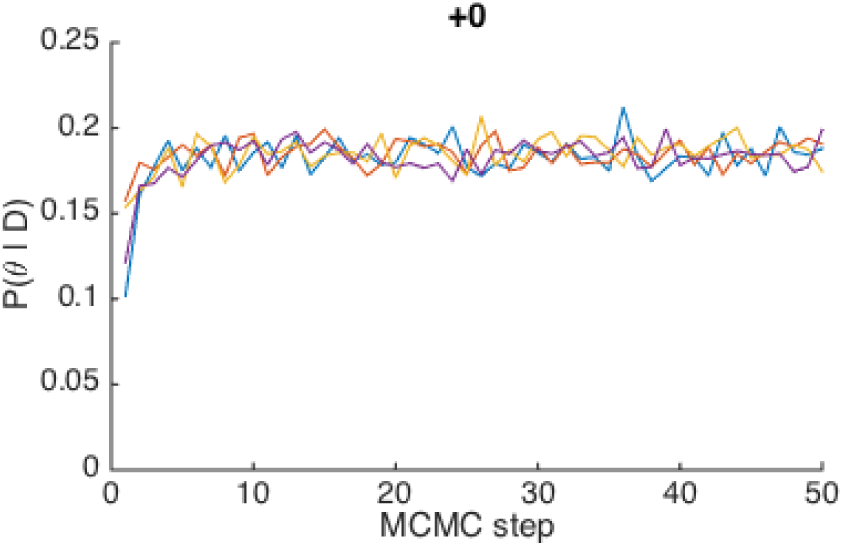
Inferred *θ*_+0_ at each step in four MCMC chains for +0 production

Figure 2 shows a remarkable difference between the probability of the productions that were originally used based on geometricalintuitions and the ad-hoc productions. The plot also shows that each clockwise production has almost the same probability than its corresponding anticlockwise production, and a similar relation appears between horizontal and vertical symmetry (H and V) and symmetries around diagonal axes (A and B). This is important because the original experiment was designed to balance such behavior; the inferred grammar reflects this.

Figure 4 shows the same inferred *θ* but grouped <u>accordin</u>g to production family. Grouping stresses the low probability of all the ad-hoc productions, but also shows an important difference between REP and the rest of the productions, particularly the simple concatenation of productions (CONCAT). This indicates that the *language of geometry* is capable of reusing simpler structures that capture geometrical meaning to explain the observed data, a key aspect of a successful model of LoT.

**Figure 4.**
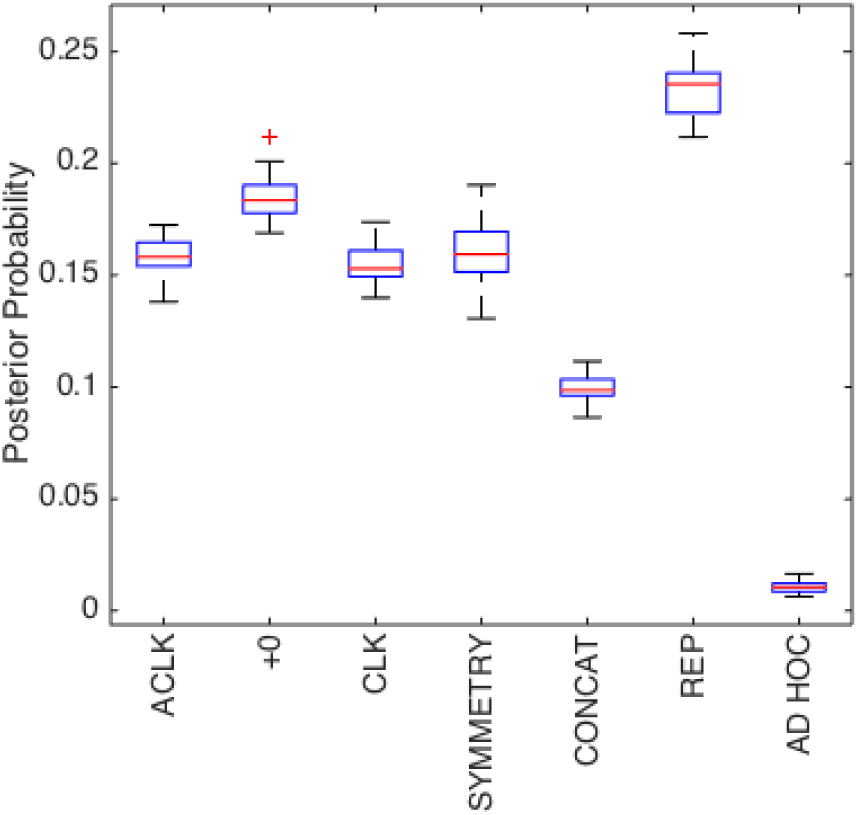
Inferred *θ*_*i*_ probability for each production in the grammar grouped by family

We then ran the same inference method using observed sequences from other experiments but only with the original grammar productions (i.e. setting aside the ad-hoc productions). We compared the result of inferring over our previously analyzed sequences generated by adults with sequences generated by children (experiment 2 from Amalric et al. (2017)) and the actual expected sequences for an ideal player.

Figure 5 shows the probabilities for each atomic production that is inferred after each population. The figure denotes that different populations can converge to different probabilities and thus different LoTs. Specifically, it is worth mentioning that the ideal learner indeed uses more repetition productions than simple concatenations when compared to adults. In the same way, adults use more repetitions than children. This could mean that the ideal learner is capable of reproducing the sequences by recursively embedding other smaller programs, whereas adults and children more so have problems understanding or learning the smaller concept that can explain all the sequences from the experiments, which is consistent with the results from the MDL model in (Amalric et al., 2017).

**Figure 5.**
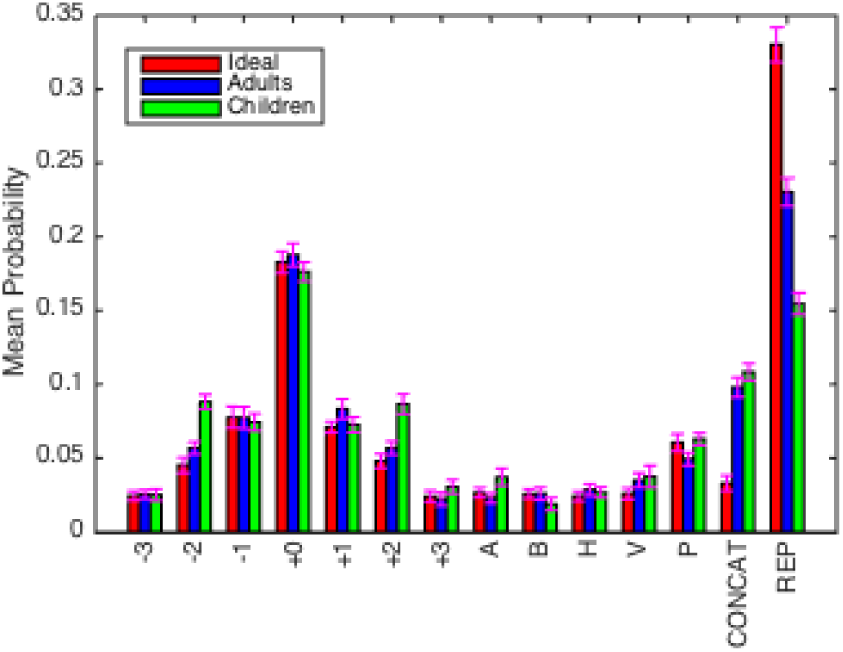
Inferred *θ*_*i*_ for Ideal learner, Adults and Children

It is worth mentioning that in (Amalric et al., 2017) the complete grammar for the *language of geometry* could explain adults’ behavior but had problems to reproduce the children’s patterns for some sequences. However, they also showed that a reduced set of productions that penalizes the rotational symmetry (P) could adequately explain children’s behavior. In Figure 5 we do not see any significant difference for this production between children and adults. This might not necessarily be contradictory, as the model for children in (Amalric et al., 2017) used a MDL approach for composing productions that took into account the occurrences of the rotational symmetry in the *minimal* program of each sequence. On the other hand, the Bayesian model in this work tries to explain the observed sequences considering the probability of a sequence summing over *all* the possible programs that can generate it. Thus, a production that is not part of the minimal program for a sequence might not necessarily be less probable when considering the entire distribution of programs for that same sequence.

## Coding Theorem

For each phenomenon there can always be an extremely large, possible infinite, number of explanations. In a LoT model, this space is constrained by the grammar *𝒢* that defines the valid hypotheses in the language. Still, one has to define how a hypothesis is chosen among all possibilities. Occam’s razor says that amongst all possible hypothesis that explain a phenomenon, one should choose the simplest. In cognitive science, the MDL framework has been widely used to model such bias in human cognition, and in *the language of geometry* in particular Amalric et al. (2017). The MDL framework is based on the ideas of information theory (Shannon, 1948), Kolmogorov complexity (Kolmogorov, 1968) and Solomonoff induction (Solomonoff, 1964).

Occam’s razor was formalized by Solomonoff (1964) in his theory of universal inductive inference, which proposes a universal prediction method that successfully approximates any distribution *μ* based on previous observations, with the only assumption of *μ* being computable. In short, Solomonoff’s theory uses all programs (in the form of prefix Turing machines) that can describe previous observations of a sequence to calculate the probability of the next symbols in an optimal fashion, giving more weight to shorter programs. Intuitively, simpler theories with low complexity have higher probability than theories with higher complexity. Formally, this relationship is described by the Coding Theorem (Levin, 1974), which closes the gap between the concepts of Kolmogorov complexity and probability theory. However, LoT models that define a probabilistic distribution for their hypotheses do not attempt to compare it with a complexity measure of the hypotheses like the ones used in MDL, nor the other way around.

In what follows we formalize the Coding Theorem (for more information, see (Li & Vitányi, 2013)) and test it experimentally. To the best our knowledge, this is the first attempt to validate these ideas for a particular (non universal) language. The reader should note that we are not validating the theorem itself as it has already been proved for universal Turing Machines. Here, we are testing whether the inverse logarithmic relationship between the probability and complexity holds true when defined for a specific non universal language.

## The formal statement

Let *M* be a prefix Turing machine –by *prefix* we mean that if *M*(*x*) is defined, then *M* is undefined for every proper extension of *x*. Let *P*_*M*_(*x*) be the probability that the machine *M* computes output *x* when the input is filled-up with the results of fair coin tosses, and let *K*_*M*_(*x*) be the *Kolmogorov complexity of x relative to M*, which is defined as the length of the shortest program which outputs *x*, when executed on *M*. The Coding Theorem states that for every string *x* we have

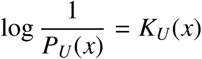

up to an additive constant, whenever *U* is a *universal* prefix Turing machine –by *universal* we mean a machine which is capable of simulating every other Turing machine; it can be understood as the underlying (Turing-complete) chosen programming language. It is important to remark that neither *P*_*U*_, nor *K*_*U*_ are computable, which means that such mappings cannot be obtained through effective means. However, for specific (non-universal) machines *M*, one can, indeed, compute both *P*_*M*_ and *K*_*M*_.

## Testing the Coding Theorem for *𝒢 eo*

Despite the fact that *P*_*M*_ and *K*_*M*_ are defined over a Turing Machine *M*, the reader should note that a LoT is not usually formalized with a Turing Machine, but instead as a programming language with its own syntax of valid programs and semantics of execution, which stipulates how to compute a concept from a program. However, one can understand programming languages as defining an equivalent (not necessarily universal) Turing Machine model, and a LoT as defining its equivalent (not necessarily universal) Turing Machine *𝒢*. In short, machines and languages are interchangeable in this context: they both specify the programs/terms, which are symbolic objects that, in turn, describe semantic objects, namely, strings.

### The Kolmogorov complexity relative to 𝒢eo

In (Amalric et al., 2017), the Minimal Description Length was used to model the combination of productions from the *language of geometry* into concepts by defining a Kolmogorov complexity relative to the *language of geometry*, which we denote *K* _*𝒢 eo*_. *K* _*𝒢 eo*_(*x*) is the minimal size of an expression in the grammar of *𝒢eo* which describes *x*. The formal definition of ‘size’ can be found in the cited work but in short: each of the atomic productions adds a fixed cost of 2 units; using any of the repetition productions to iterate *n* times a list of other productions adds the cost of the list, plus ⌊log(*n*)⌋; and joining two lists with a concatenation costs the same as the sum of the costs of both lists.

### The probability relative to 𝒢eo

On the other hand, with the Bayesian model specified in this work, we can define *P*(*x* | *𝒢eo*, *θ*) which is the probability of a string *x* relative to *𝒢eo* and its vector of probabilities for each of the productions.

For the sake of simplicity, we will use *P*_*𝒢eo*_(*x*) to denote *P*(*x* | *𝒢eo*, *θ*) when *θ* is the inferred probability from the observed adult sequences from the experiment.

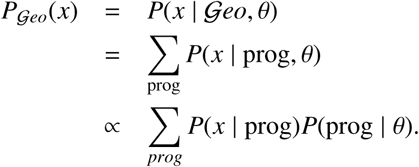

Here, we calculate both *P*_*𝒢eo*_ (*x*) and *K*_*𝒢eo*(*x*)_ in an exact way (note that *𝒢eo*, seen as a programming language, is not Turing-complete). In this section, we show an experimental equivalence between such measures which is consistent with the Coding Theorem. We should stress, once more, that the theorem does not predict that this relationship should hold for a specific non-universal Turing Machine.

To calculate *P*_*𝒢eo*_(*x*) we are not interested in the normalization factor of *P*(*x* | prog)*P*(prog | *θ*) because we are just trying to measure the relationship between *P*_*𝒢eo*_ and *K*_*𝒢eo*_ in terms of the Coding Theorem. Note, however, that calculating *P*_*𝒢eo*_(*x*) involves calculating all programs that compute each of the sequences as in our previous experiment. To make this tractable we calculated *P*_*𝒢eo*_(*x*) for 10,000 unique random sequences for each of the possible sequence lengths from the experiment (i.e., up to eight). When the length of the sequence did not allow 10,000 unique combinations, we used all the possible sequences of that length.

## Coding Theorem Results

Figure 6 shows the mean probability *P*_*𝒢eo*_(*x*) for all sequences *x* with the same value of *K*_*𝒢eo*(*x*)_ and length between 4 and 8 (|*x*| ∈ [4, 8]) for all generated sequences *x*. The data is plotted with a logarithmic scale for the x-axis, illustrating the inverse logarithmic relationship between *K*_*𝒢eo*_(*x*) and *P*_*𝒢eo*_(*x*). The fit is very good, with *R*^2^ = .99, in *R*^2^ = .94, *R*^2^ = .97, *R*^2^ = .99 and *R*^2^ = .98 for Figure 6a, Figure 6b, Figure 6c, Figure 6d and Figure 6e, respectively.

**Figure 6.**
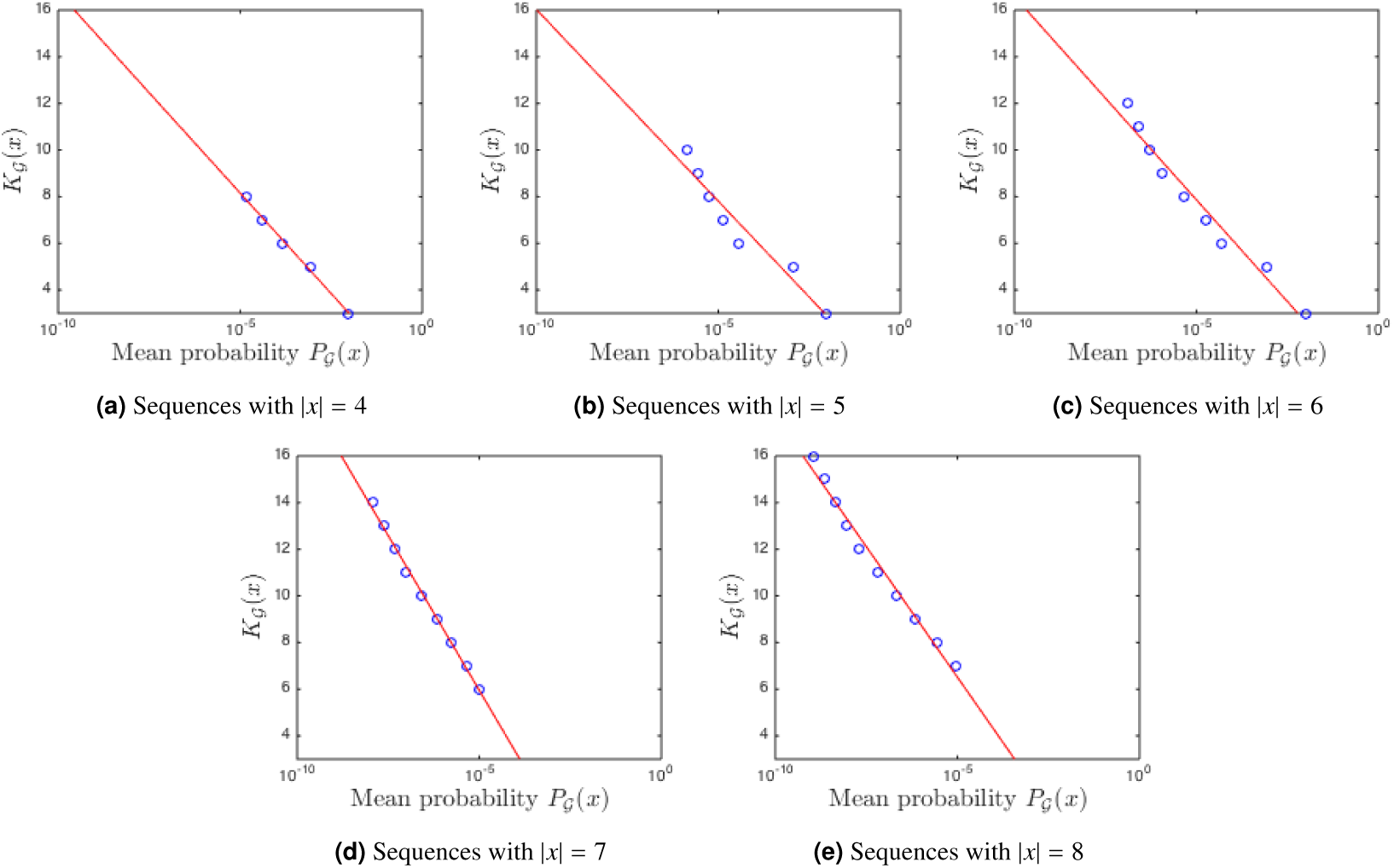
Mean probability *P*_*𝒢eo*_(*x*) for all sequences *x* with the same complexity

This relationship between the complexity *K*_*𝒢eo*_ and the probability *P*_*𝒢eo*_ defined for finite sequences in the *language of geometry*, matches the theoretical prediction for infinite sequences in universal languages described in the Coding Theorem. At the same time, it captures the Occam’s razor intuition that the simpler sequences one can produce or explain with this language are also the more probable.

Figures 7 and 8 show the histogram of *P*_*𝒢eo*_(*x*) and *K*_*𝒢eo*_(*x*), respectively, for sequences with length = 8 to get a better insight about both measures. The histogram of the rest of the sequence’s lengths are included in Figures B1 and B2 for completeness, and they all show the same behavior.

**Figure 7.**
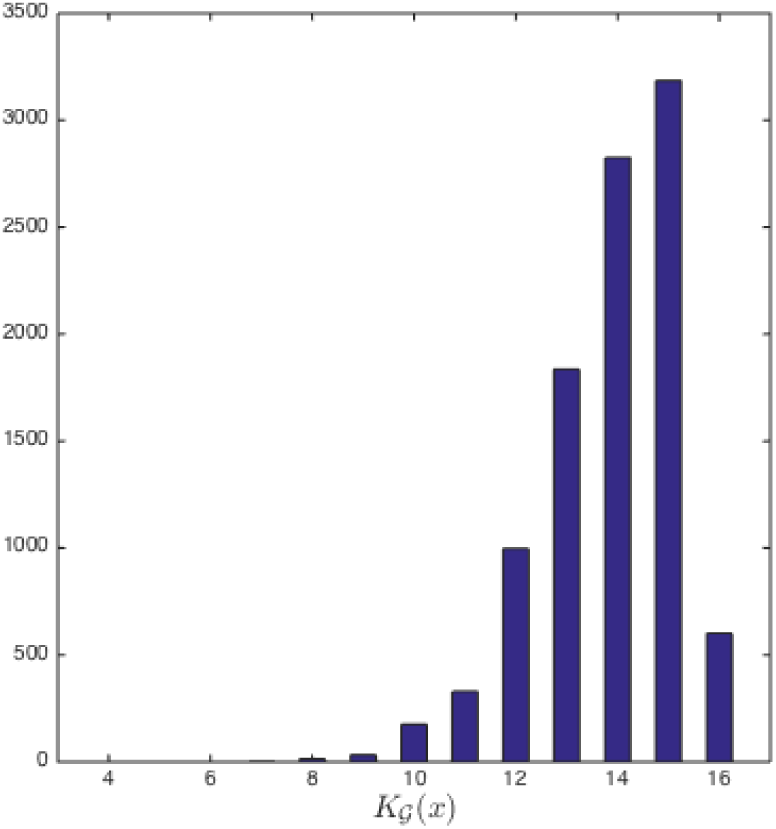
Histogram of complexity *K*_*𝒢eo*_(*x*) for sequences *x* with |*x*| = 8

**Figure 8.**
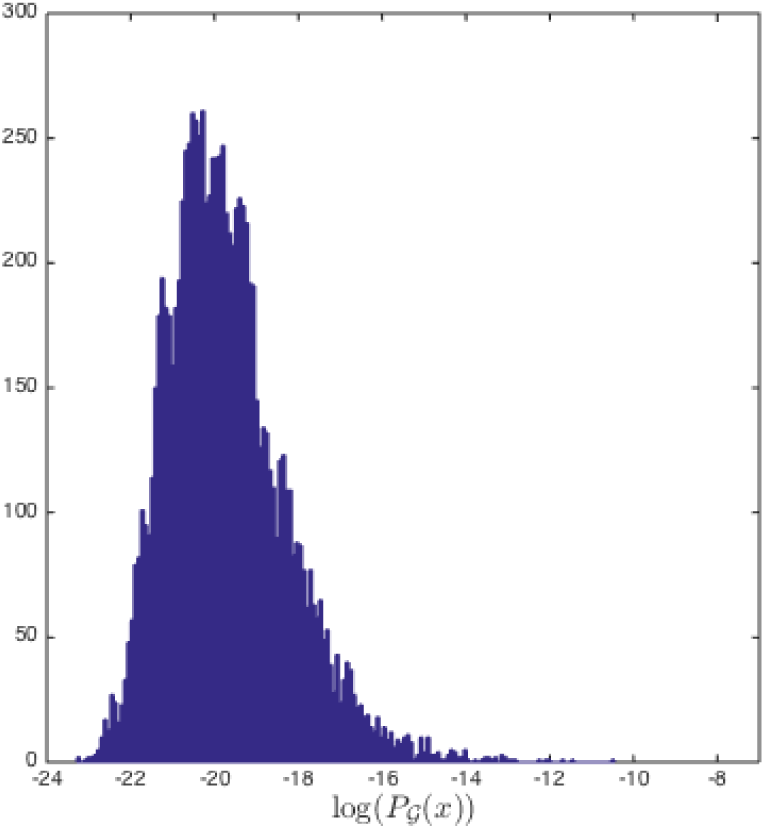
Histogram of probability *P*_𝒢*eo*_(*x*) for sequences *x* with |*x*| = 8

## Discussion

We have presented a Bayesian inference method to select the set of productions for a LoT and test its effectiveness in the domain of a geometrical cognition task. We have shown that this method is useful to distinguish between arbitrary ad-hoc productions and productions that were intuitively selected to mimic human abilities in such domain.

The proposal to use Bayesian models tied to PCFG grammars in a LoT is not new. However, previous work has not used the inferred probabilities to gain more insight about the grammar definition in order to modify it. Instead, it had usually integrated out the production probabilities to better predict the data, and even found that hierarchical priors for grammar productions show no significant differences in prediction results over uniform priors (Piantadosi et al., 2012; Yildirim & Jacobs, 2015).

We believe that inferring production probabilities can help prove the adequacy of the grammar, and can further lead to a formal mechanism for selecting the correct set of productions when it is not clear what a proper set should be. Researchers could use a much broader set of productions than what might seem intuitive or relevant for the domain and let the hierarchical Bayesian inference framework select the best subset.

Selecting a broader set of productions still leaves some arbitrary decisions to be made. However, it can help to build a more robust methodology that – combined with other ideas like testing grammars with different productions for the same task (Piantadosi et al., 2016)– could provide more evidence of the adequacy of the proposed LoT.

Having a principled method for defining grammars in LoTs is a crucial aspect for their success because slightly different grammars can lead to different results, as has been shown in (Piantadosi et al., 2016).

The experimental data used in this work was designed by Amalric et al. (2017) to understand how humans encode visuo-spatial sequences as structured expressions. As future research, we plan to perform a specific experiment to test these ideas in a broader range of domains. Additionally, data from more domains is needed to demonstrate if this method could also be used to effectively prove whether different people use different LoT productions as outlined in Figure 5.

Finally, we showed an empirical equivalence between the complexity of a sequence in a minimal description length (MDL) model and the probability of the same sequence in a Bayesian inference model which is consistent with the theoretical relationship described in the Coding Theorem. This opens an opportunity to bridge the gap between these two approaches that had been described ad complementary by some authors MacKay (2003, chapter 28.3).

## Appendix A

MCMC steps for the rest of *𝒢eo*’s grammar productions

**Figure A1.**
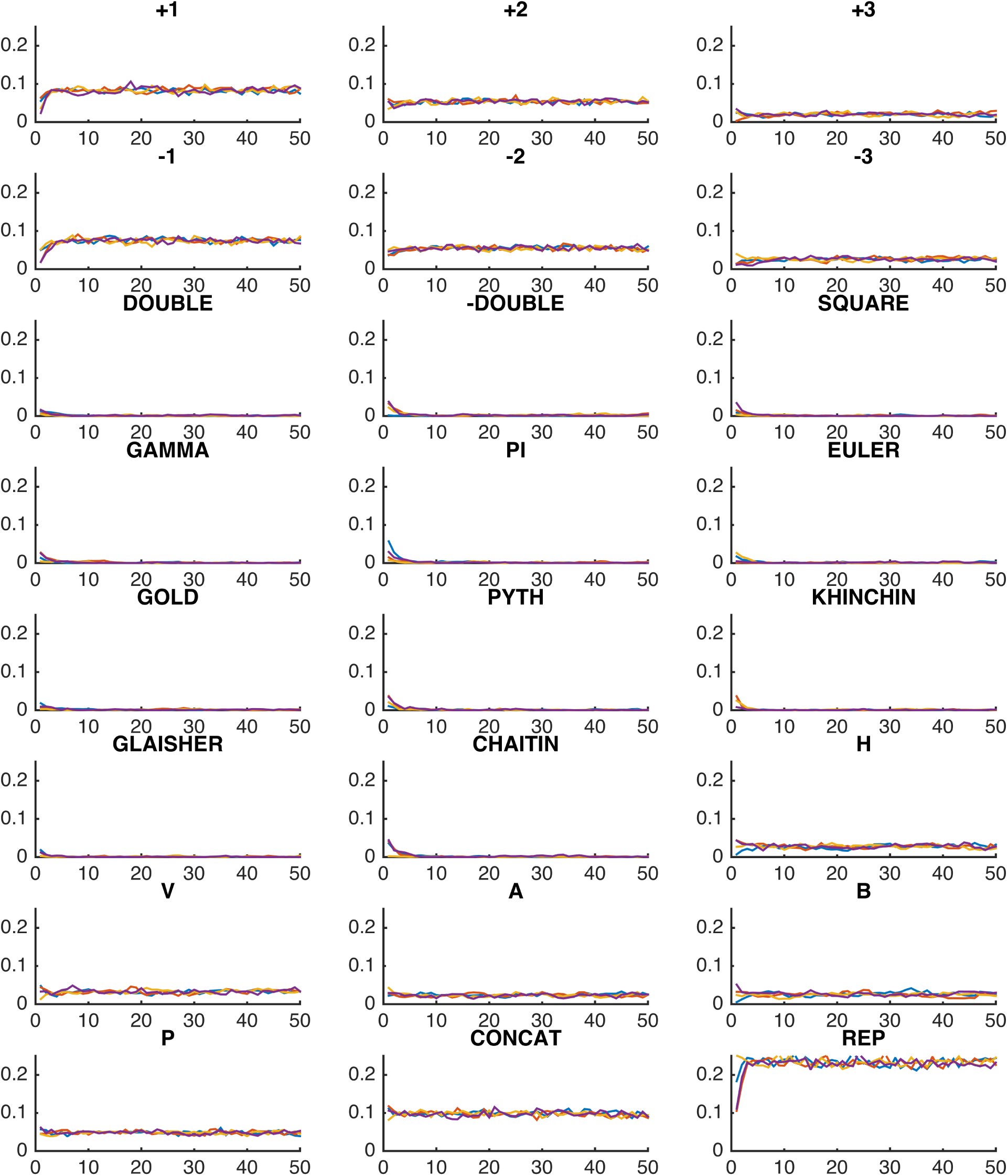
Inferred *θ*_*i*_ at each step in four MCMC chains

## Appendix B

Histograms of *P*_*𝒢eo*_(*x*) and *K*_*𝒢eo*_(*x*)

**Figure B1.**
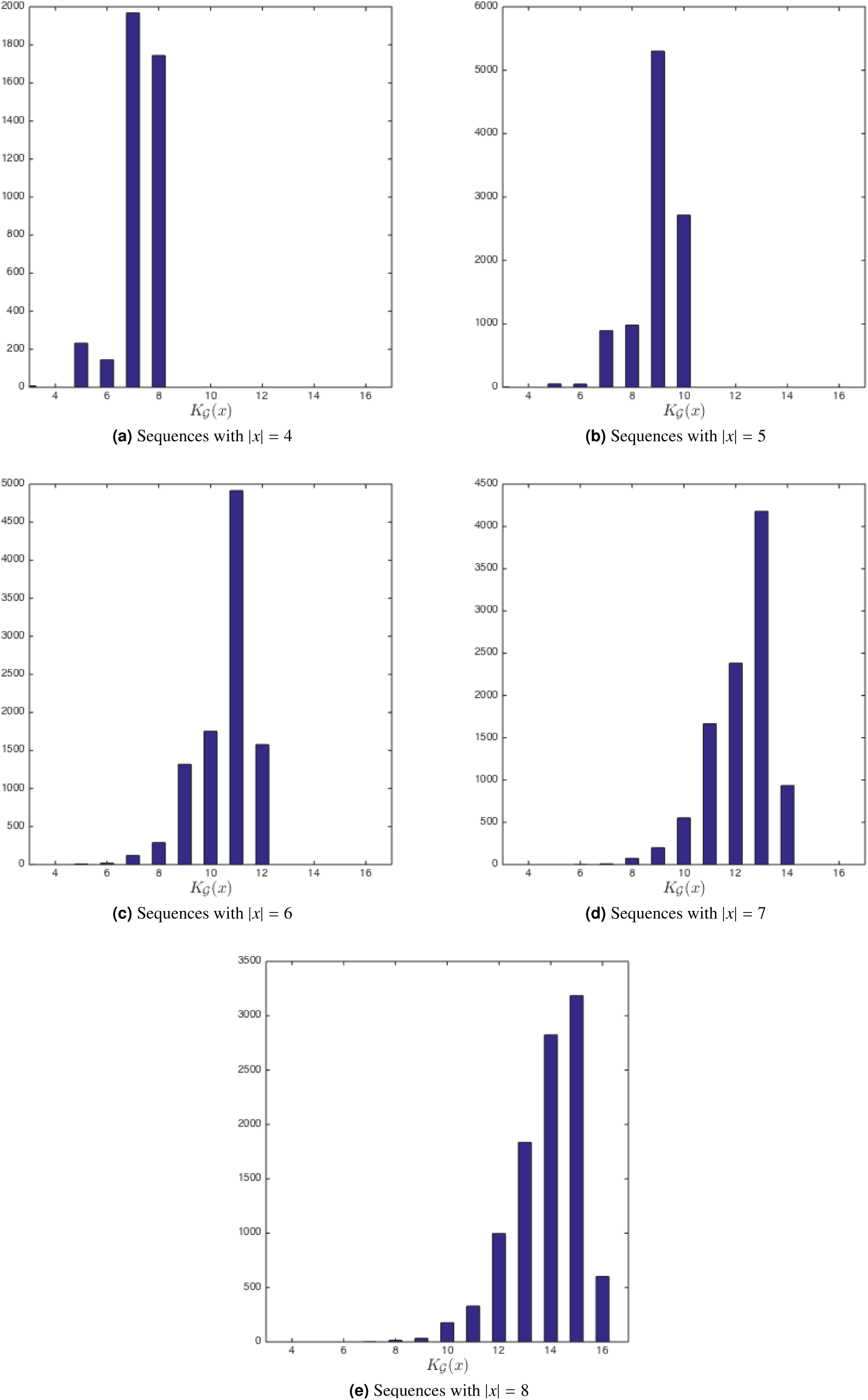
Histogram of complexity *K*_*𝒢eo*_(*x*) for sequences *x*

**Figure B2.**
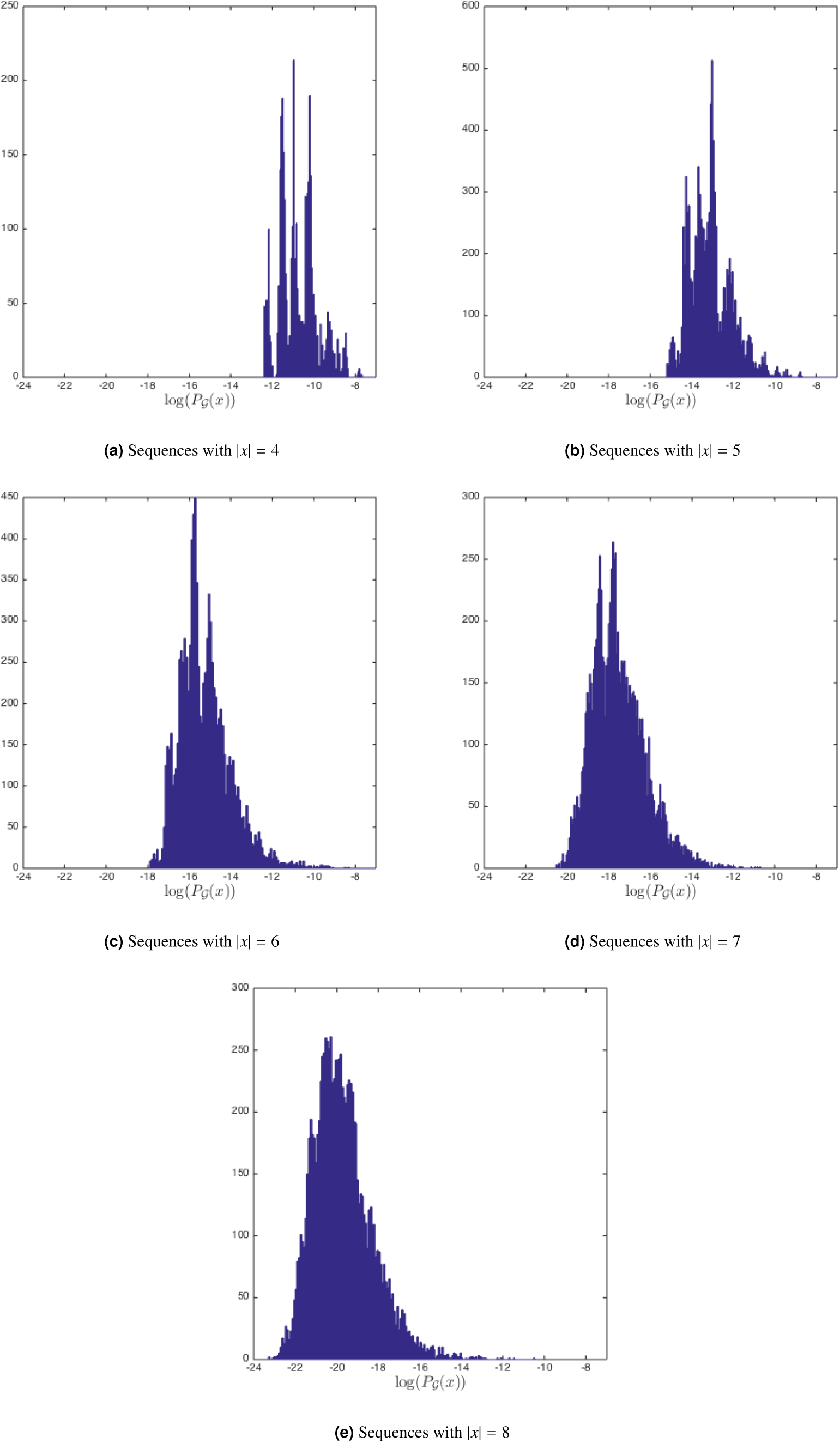
Histogram of probability *P*_*𝒢eo*_(*x*) for sequences *x*

## Footnotes

1 Calculated for a particular universal Turing Machine and programs up to 84 bits long (Calude, Dinneen, Shu, et al., 2002)

## References

Amalric, M., Wang, L., Pica, P., Figueira, S., Sigman, M., & Dehaene, S. (2017). The language of geometry: Fast comprehension of adults and preschoolers. PLOS Computational Biology, 13(1), e1005273.

Aydede, M. (1997). Language of thought: The connectionist contribution. Minds and Machines, 7(1), 57–101.

Blackburn, S. (1984). *Spreading the word:* Grounding in the philosophy of language. Clarendon Press.

Boole, G. (1854). *An investigation of the laws of thought: on which are founded the mathematical theories of logic and probabilities*. Dover Publications.

Borges, J. L. (1944). Ficciones, 1935-1944. Buenos Aires: Sur.

Calude, C. S., Dinneen, M. J., Shu, C.-K., et al. (2002). Computing a glimpse of randomness. Experimental Mathematics, 11(3), 361–370.

Dehaene, S., Izard, V., Pica, P., & Spelke, E.(2006). Core knowledge of geometry in an amazonian indigene group. Science,311(5759), 381–384.

Dillon, M. R., Huang, Y., & Spelke, E. S. (2013). Core foundations of abstract geometry. Proceedings of the National Academy of Sciences, 110(35), 14191–14195.

Ellis, K., Solar-Lezama, A., & Tenenbaum, J.(2015). Unsupervised learning by program synthesis. In Advances in neural information processing systems (pp. 973–981).

Fodor, J. (1975). *The language of thought*. Harvard University Press.

Geman, S., & Geman, D. (1984). Stochastic relaxation, gibbs distributions, and the bayesian restoration of images. Pattern Analysis and Machine Intelligence, IEEE Transactions on(6), 721–741.

Gentner, D. (1983). Structure-mapping: A theoretical framework for analogy. Cognitive science, 7(2), 155–170.

Goldsmith, J. (2001). Unsupervised learning of the morphology of a natural language. Computational linguistics, 27(2), 153–198.

Goldsmith, J. (2002). Probabilistic models of grammar: Phonology as information minimization. Phonological Studies, 5, 21–46.

Goodman, N. D., Tenenbaum, J. B., Feldman, J., & Griffiths, T. L. (2008). A rational analysis of rule-based concept learning. Cognitive Science, 32(1), 108–154.

Izard, V., Pica, P., Dehaene, S., Hinchey, D., & Spelke, E. (2011). Geometry as a universal mental construction. Space, Time and Number in the Brain, 19, 319–332.

Johnson, M., Griffiths, T. L., & Goldwater, S.(2007). Bayesian inference for pcfgs via markov chain monte carlo. In Hlt-naacl (pp. 139–146).

Knowles, J. (1998). The language of thought and natural language understanding. Analysis, 58(4), 264–272.

Kolmogorov, A. N. (1968). Three approaches to the quantitative definition of information*. International Journal of Computer Mathematics, 2(1-4), 157–168.

Landau, B., Gleitman, H., & Spelke, E. (1981). Spatial knowledge and geometric representation in a child blind from birth. Science, 213(4513), 1275–1278.

Lee, S. A., Sovrano, V. A., & Spelke, E. S. (2012). Navigation as a source of geometric knowledge: Young children's use of length, angle, distance, and direction in a reorientation task. Cognition, 123(1), 144–161.

Levin, L. A. (1974). Laws of information conservation (nongrowth) and aspects of the foundation of probability theory. Problemy Peredachi Informatsii, 10(3), 30–35.

Li, M., & Vitányi, P. (2013). An introduction to kolmogorov complexity and its applications. Springer Science & Business Media.

Loewer, B., & Rey, G. (1991). Meaning in mind. Fodor and his Critics.

Machilsen, B., Pauwels, M., & Wagemans, J. (2009). The role of vertical mirror symmetry in visual shape detection. Journal of Vision, 9(12), 11–11.

MacKay, D. J. (2003). Information theory, inference and learning algorithms. Cambridge university press.

Manning, C., & Schütze, H. (1999). Foundations of statistical natural language processing. MIT Press.

Nosofsky, R. M. (1986). Attention, similarity, and the identification–categorization relationship. Journal of experimental psychology: General, 115(1), 39.

Piantadosi, S. T., & Jacobs, R. A. (2016). Four problems solved by the probabilistic language of thought. Current Directions in Psychological Science, 25(1), 54–59.

Piantadosi, S. T., Tenenbaum, J. B., & Goodman, N.D. (2012). Bootstrapping in a language of thought: A formal model of numerical concept learning. Cognition, 123(2), 199–217.

Piantadosi, S. T., Tenenbaum, J. B., & Goodman, N. D. (2016). The logical primitives of thought: Empirical foundations for compositional cognitive models.

Romano, S., Sigman, M., & Figueira, S. (2013).: A language of thought with turing-computable kolmogorov complexity. Papers in Physics, 5, 050001.

Rosch, E. (1999). Principles of categorization. Concepts: core readings, 189.

Rosch, E., & Mervis, C. B. (1975). Family resemblances: Studies in the internal structure of categories. Cognitive psychology, 7(4), 573–605.

Rosch, E., Simpson, C., & Miller, R. S. (1976). Structural bases of typicality effects. Journal of Experimental Psychology: Human perception and performance, 2(4), 491

Shannon, C. (1948). A mathematical theory of communication. Bell System Technical Journal, 27, 379–423, 623–656.

Solomonoff, R. J. (1964). A formal theory of inductive inference. part i. Information and control, 7(1), 1–22.

Tenenbaum, J. B., Kemp, C., Griffiths, T. L., & Goodman, N. D. (2011). How to grow a mind: Statistics, structure, and abstraction. science, 331(6022), 1279–1285.

Ullman, T. D., Goodman, N. D., & Tenenbaum, J. B. (2012). Theory learning as stochastic search in the language of thought. Cognitive Development, 27(4), 455–480.

Westphal-Fitch, G., Huber, L., Gómez, J. C., & Fitch, W. T. (2012). Production and perception rules underlying visual patterns: effects of symmetry and hierarchy. Philosophical Transactions of the Royal Society of London B: Biological Sciences, 367(1598), 2007–2022.

Yildirim, I., & Jacobs, R. A. (2015). Learning multisensory representations for auditory-visual transfer of sequence category knowledge: a probabilistic language of thought approach. Psychonomic bulletin & review, 22(3), 673–686.

